# Metagenomic Insights Reveal Unrecognized Diversity of Entotheonella in Japanese *Theonella* Sponges

**DOI:** 10.1101/2024.01.29.577764

**Authors:** Sota Yamabe, Kazutoshi Yoshitake, Akihiro Ninomiya, Jörn Piel, Haruko Takeyama, Shigeki Matsunaga, Kentaro Takada

**Author notes:** Corresponding author Kentaro Takada. Jörn Piel.

## Abstract

Numerous biologically active natural products have been discovered from marine sponges, particularly from *Theonella swinhoei*, which is known to be a prolific source of natural products such as polyketides and peptides. Recent studies have revealed that many of these natural products are biosynthesized by *Candidatus* Entotheonella phylotypes, which are uncultivated symbionts within *T. swinhoei*. Consequently, Entotheonella is considered an untapped biochemical resource. In this study, we conducted metagenomic analyses to assess the diversity of Entotheonella in four *T. swinhoei* individuals (two each of chemotypes Y and W), after separating filamentous bacteria using density gradient centrifugation. We obtained five Entotheonella genomes from filamentous bacteria-enriched fractions. Notably, one of these genomes is significantly different from previously reported Entotheonella variants. Additionally, we identified closely related Entotheonella members present across different chemotypes of *T. swinhoei*. Thus, our metagenomic insights reveal that the diversity of Entotheonella within *Theonella* sponges is greater than previously recognized.

## Introduction

Marine sponges, representing one of the most ancient forms of multicellular life, are fascinating not only in their own rights but also serve as critical models for understanding primitive animal-microbe symbiosis. The complex microbiomes they harbor, comprising various microorganisms, plays a pivotal role in their survival and offer significant insights into the nature of symbiotic relationships (Hentschel et al. 2002; Taylor et al. 2007). Previous research has shown that symbiotic microorganisms can occupy up to 40% of the sponge volume (Taylor et al. 2007) and are believed to significantly contribute to the sponge’s chemical defense and overall health (Bewley et al. 1998; Piel et al. 2004). Despite their importance, detailed biological information, including genomes and life cycles of most symbiotic microorganisms, remains elusive due to their uncultured status.

The marine sponge *Theonella swinhoei*, with its distinct Yellow (TSY) and White (TSW) chemotype, present a unique model for studying sponge-microbes interaction in terms of secondary metabolites. TSY and TSW, being found throughout the Western Pacific and the Red Sea, have been a source of over 150 natural products with diverse biological activities and chemical characteristics (Bewley et al. 1998; Winder et al. 2011; Bewley et al. 1996; Hamada et al. 2005). Interestingly, there is minimal overlap in the metabolites found in the two chemotypes (Wegerski et al. 2007), which makes *Theonella* sponges attractive bioresources for drug lead discovery. For a long time, the actual producers of sponge-derived bioactive metabolites were unknown. The discovery of the antifungal peptides, theopalauamide in the filamentous bacteria-enriched fraction of *T. swinhoei* in Palau marked a significant breakthrough (Bewley et al. 1996; Schmidt et al. 1998). This finding implied that a filamentous bacterium, named *Candidatus* Entotheonella palauensis (Schmidt et al. 2000), is the producer of theopalauamide. Further research genetically revealed that *Candidatus* Entotheonella factor is responsible for the production of almost all of the polyketides and peptides found in TSY, such as onnamides, keramamides, nazumamides, cyclotheoneamides, konbamides, and polytheonamides (Wilson et al. 2014). Recent research on the TSW metagenome suggests that *Candidatus* Entotheonella serta is a producer of theonellamides (Mori et al. 2018) and misakinolide. Thus, Entotheonella represents a taxon with broad symbiosis in *Theonella* sponges and an exceptional ability to produce secondary metabolites.

In this study, we aim to explore the further diversity of Entotheonella in the fractionated symbiotic microorganisms derived from Japanese TSY and TSW using density gradient centrifugation. This process yielded three and five Entotheonella-enriched fractions from TSY and TSW, respectively. Notably, the morphology of Entotheonella is distinctly different across these fractions, prompting us to conduct a comprehensive metagenomic analysis. Through this study, we provide further insight into the diversity of Entotheonella in terms of genome content and secondary metabolites.

## Materials and Methods

### Sponge Collection, Preparation of Entotheonella-Enriched Fraction, and Metagenome Analyses

Sponge specimens, *Theonella swinohoei* YA, YB, WC, and WD, were collected by scuba diving near Hachijo-jima, Japan (33° 1437 N, 139° 7996 E) during 2021-2022. After crushing the sponges, the samples were filtered through a 50 µm mesh and suspended in Ca^2+^- and Mg^2+^-free seawater to disperse intercellular aggregations. For isolating filamentous bacteria-enriched fractions, density gradient centrifugation was employed. Percoll was diluted to 50%, 40%, 30%, and 20% with seawater containing 18 mM 2-(*N*-morpholino) ethanesulfonic acid (MES). These solutions were then layered in centrifuge tubes in descending order of dilution, and pre-centrifuged at 10,000 rpm for 60 minutes to establish a precise density gradient. The sponge sample suspended in seawater was added to this gradient, and the filamentous bacteria-enriched fractions were collected after centrifugation at 3,000 rpm for 8 min. The filamentous bacteria fractions, diluted with 38% iodixanol (IDX) in seawater containing 20 mM 4-(2-hydroxyethyl)-1-piperazine ethanesulfonic acid (HEPES), was added to centrifuge tubes and then layered in descending order of dilution at 36%, 34%, 32%, 30%, and 26% IDX solutions. Genomic DNA was extracted using the phenol-chloroform method. The lysis buffer for DNA extraction was composed of 50 mM Tris-HCl (pH 7.5), 50 mM EDTA, 350 mM NaCl, 2% SDS, and 8 M urea. A volume of lysis buffer ten times the packed volume of the collected filamentous bacteria-enriched fractions was used. The mixture was incubated at 60 °C for 120 min, followed by centrifugation at 500 g for 5 min. DNA was then extracted with equal volumes of phenol/chloroform/isoamyl alcohol (25:24:1) twice, and chloroform/isoamyl alcohol (24:1) three times, using the collected supernatant. The DNA was precipitated by adding 0.1x volume of 3 mM sodium acetate (pH 5.2) and 2.5x volume of 99.5% cold ethanol, and subsequently dissolved in Tris-EDTA buffer. Metagenomic sequencing was performed using the Illumina HiSeq X platform.

### Bioinformatics Analyses

De novo assembly of shotgun sequencing data was conducted using the CLC workbench to obtain draft sequences. Sequence coverage was calculated using bbmap, and metagenomic binning, based on tetranucleotide frequency and coverage, was performed with MetaBAT2 (Bushnel. 2015, Kang et al. 2019). Contig clusters closely related to Entotheonella were identified by comparing them to known genomes. The binning results were visualized in R using the gbtools package (Seah et al. 2015). The draft genomes resembling Entotheonella were then evaluated for completeness and contamination using CheckM2 (Alex et al. 2022), which assesses genome quality based on single-copy gene markers. Average Nucleotide Identity (ANI) analysis, used for comparing each Entotheonella-like draft genome with established genomes, was performed using the JSpecies (Richter et al. 2016) web server tool. Additionally, a genome-level phylogenetic tree was constructed using the autoMLST (Multi Locus Species Tree) web server tool (Alanjary et al. 2019). For the identification of compound biosynthetic gene clusters (BGCs), the Antibiotics and Secondary Metabolite Analysis Shell (antiSMASH) version 7.0 (Blin et al. 2023) was utilized. When the contigs in the genomes are too fragmented, we concurrently mapped the raw reads to the known Biosynthetic Gene Clusters (BGCs) to determine whether the BGCs are present. All manual annotations and bioinformatics analyses were conducted using Geneious. Furthermore, Clinker was employed for the alignment and visualization of BGCs identified by antiSMASH (Gilchrist et al. 2021).

### LC/MS Analyses for the Sponge Extracts

A 1 mL portion of the sponge filtrate from samples YA, YB, WC, and WD was subjected to centrifugation at 15,000 g for 5 min to obtain the precipitate. Each precipitate was then extracted with MeOH, yielding crude extracts of 21.9 mg, 84.3 mg, 30.1 mg, and 20.7 mg, respectively. All obtained extracts were analyzed using high-resolution mass spectrometry (HRMS). For the standard analysis of sponge and Entotheonella extracts, solvent gradients were employed (Solvent A: H^2^O with 0.5% acetic acid; Solvent B: acetonitrile with 0.5% acetic acid), with the gradient settings for Solvent B being 10-90% for 0–20 minutes, 95% for 20–25 minutes at a flow rate of 0.2 ml/min. The analysis was conducted on a Cosmosil 2.5C^18^ MSII (2.0 i.d. x 100 mm), with the MS operated in positive ionization mode at a scan range of *m*/*z* 200–2,000.

## Results

### Acquisition of Entotheonella Genomes

The sponge homogenates derived from two individuals of TSY (YA and YB, Fig. 1A) and TSW (WC and WD, Fig. 1E) were separated using density gradient centrifugation, yielding eight filamentous bacteria-enriched fractions: Fra1 and Fra2 from YA; Fra3 from YB; Fra4, Fra5, and Fra6 from WC; Fra7 and Fra8 from WD (Fig. S1). The morphology of the bacteria in each fraction was similar to that of the previously reported Entotheonella (Fig. 1, Fig. S2) (Schmidt et al. 2000; Wilson et al. 2014). To genetically distinguish these bacteria, we performed metagenomic sequencing for each fraction. After binning the contigs based on tetra-nucleotide frequency and sequence coverage, seven Entotheonella-like genomes were identified within the clusters by referencing the known Entotheonella genome: YA-1 (8.29 Mbp) derived from *T. swinhoei* YA; YB-1 (7.36 Mbp) from *T. swinhoei* YB; WC-1 (8.23 Mbp), WC-2 (8.11 Mbp), WC-3 (6.37 Mbp) from *T. swinhoei* WC; WD-1 (7.13 Mbp), WD-2 (6.99 Mbp) from *T. swinhoei* WD (Table 1) (Dongwan et al. 2019). An analysis using CheckM, a bioinformatic tool that evaluates the quality and completeness of microbial genomes, confirmed that the genomes exhibited completeness and contamination scores of 78-95 and 6.4-10.8 (Table 1) (Alex et al. 2022). These scores suggested that cell fractionation using density gradient centrifugation enabled us to obtain Entotheonella genomes of comparable quality to those derived from single-cell genomics.

**Table 1.**
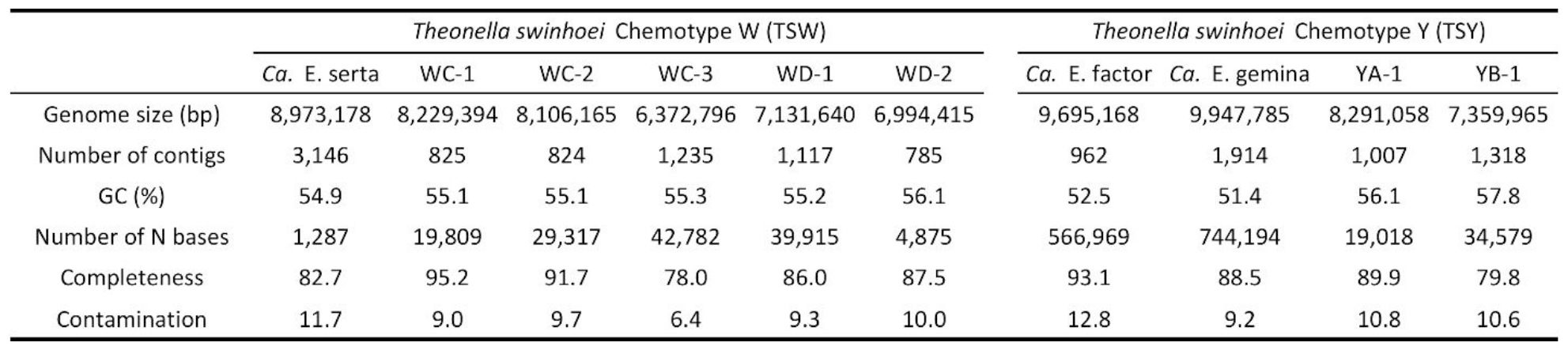
Comprehensive genomic data overview for Entotheonella phylotypes.

**Fig. 1.**
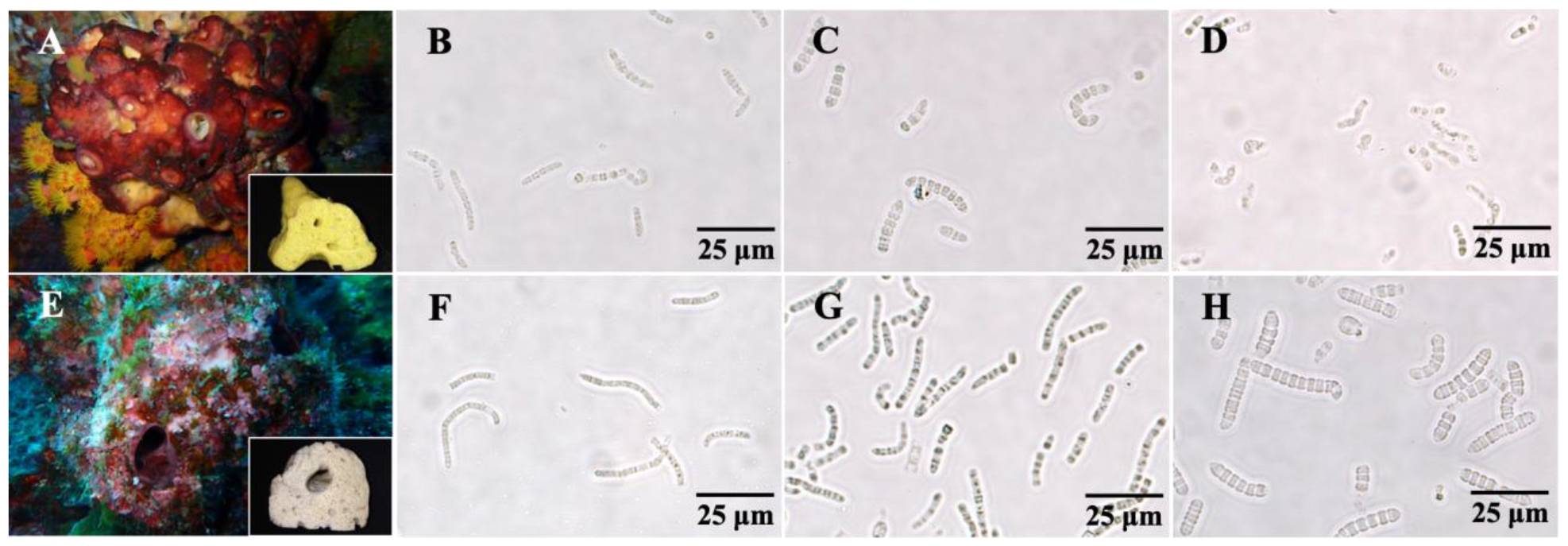
Two chemotypes of the marine sponge *Theonella swinhoei* and their associated Entotheonella-enriched fractions. (A) Shows both the external morphology and internal morphology of TSY. Micrographs depicting the filamentous bacteria-enriched fractions from (B) YA-derived Fra1, (C) Fra2 and (D) YB-derived Fra3. (E) Shows both external and internal morphology of TSW. Micrographs of the filamentous bacteria-enriched fractions from (F) WC-derived Fra4, (G) WD-derived Fra5, and (H) Fra6.

To evaluate seven Entotheonella-like genomes, we conducted an Average Nucleotide Identity (ANI) analysis (Fig. 2) (Richter et al. 2016). The analysis indicated that the acquired draft genome had 75-99 % identity with the previously reported Entotheonella. Notably, three Entotheonella genomes, WC-1, WC-2, and WC-3, showed ANI values of 98-99 % and 100% identity in their 16S rRNA genes (Fig. S3), which are identical to those of *Ca*. E. serta. The genome of WD-1 also exhibited ANI values of 96.7 % to that of *Ca*. E. serta, a score slightly above the threshold of 95-96% ANI used to define species boundaries. These data suggest that WC-1, WC-2, WC-3, and WD-1 are closely related to *Ca*. E. serta albeit with a different morphology. On the other hand, YA-1 and WD-2 exhibited ANI values of more than 99% identity in that of *Ca*. E. gemina. To our knowledge, this is the first case where the same Entotheonella variants exists in TSY and TSW, both of which differ in their chemical profiles. YB-1 showed low ANI values compared to other Entotheonella genomes, implying that it represents a novel Entotheonella candidate species (Fig. 2).

**Fig. 2.**
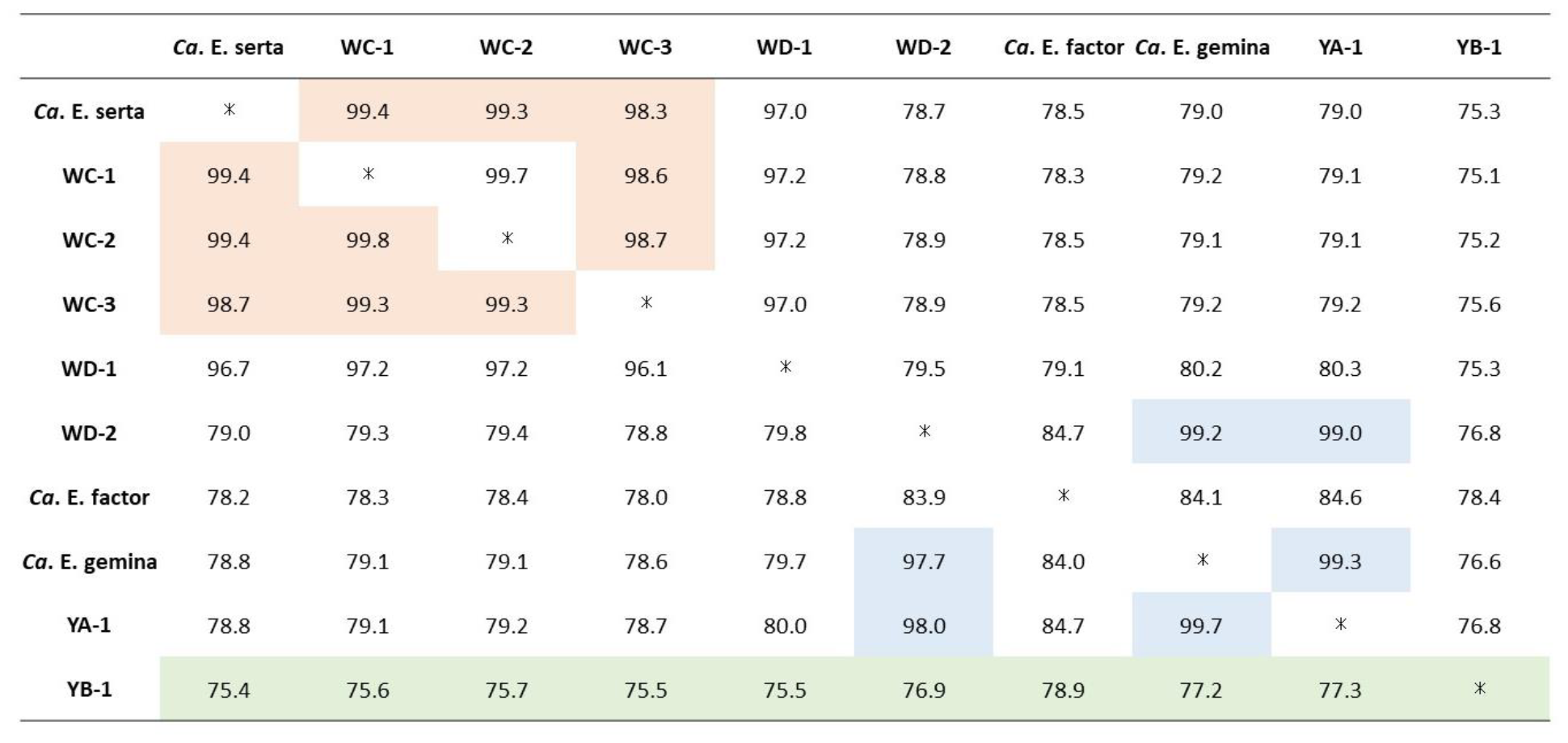
Average Nucleotide Identity (ANI) analyses for Entotheonella obtained in this study, compared with previously known phylotypes. Orange-highlighted scores indicate that WC-1, WC-2, and WC-3 are similar to *Ca*. E. serta (derived from TSW). Blue-highlighted scores indicate that YA-1 and WD-2 are similar to *Ca*. E. gemina (derived from TSY). Green-highlighted scores suggest that YB-1 represents a novel Entotheonella candidate species.

Phylogenetic classification of the species based on 16S rRNA gene sequences was challenging due to lack of sufficient sequences. Therefore, we generated a genome-level phylogenetic tree using AutoMLST, a bioinformatic tool designed for automated bacterial strain typing using whole-genome sequencing data (Table S1) (Alanjary et al. 2019). Among the five Entotheonella genomes obtained in this study and the previously known Entotheonella genome, YB-1, named *Ca*. Entotheonella arcus YB-1, formed an independent clade distinct from other Entotheonella (Fig. 3). The genome for *Ca*. E. factor, previously found in TSY, was not observed in this study. Based on the phylogenetic tree, we eventually named other phylotypes obtained in this study as *Ca*. E. gemina YA-1, *Ca*. E. gemina WD-2, *Ca*. E.serta WC-1, and *Ca*. E.serta WD-1 (Fig. 3).

**Fig. 3.**
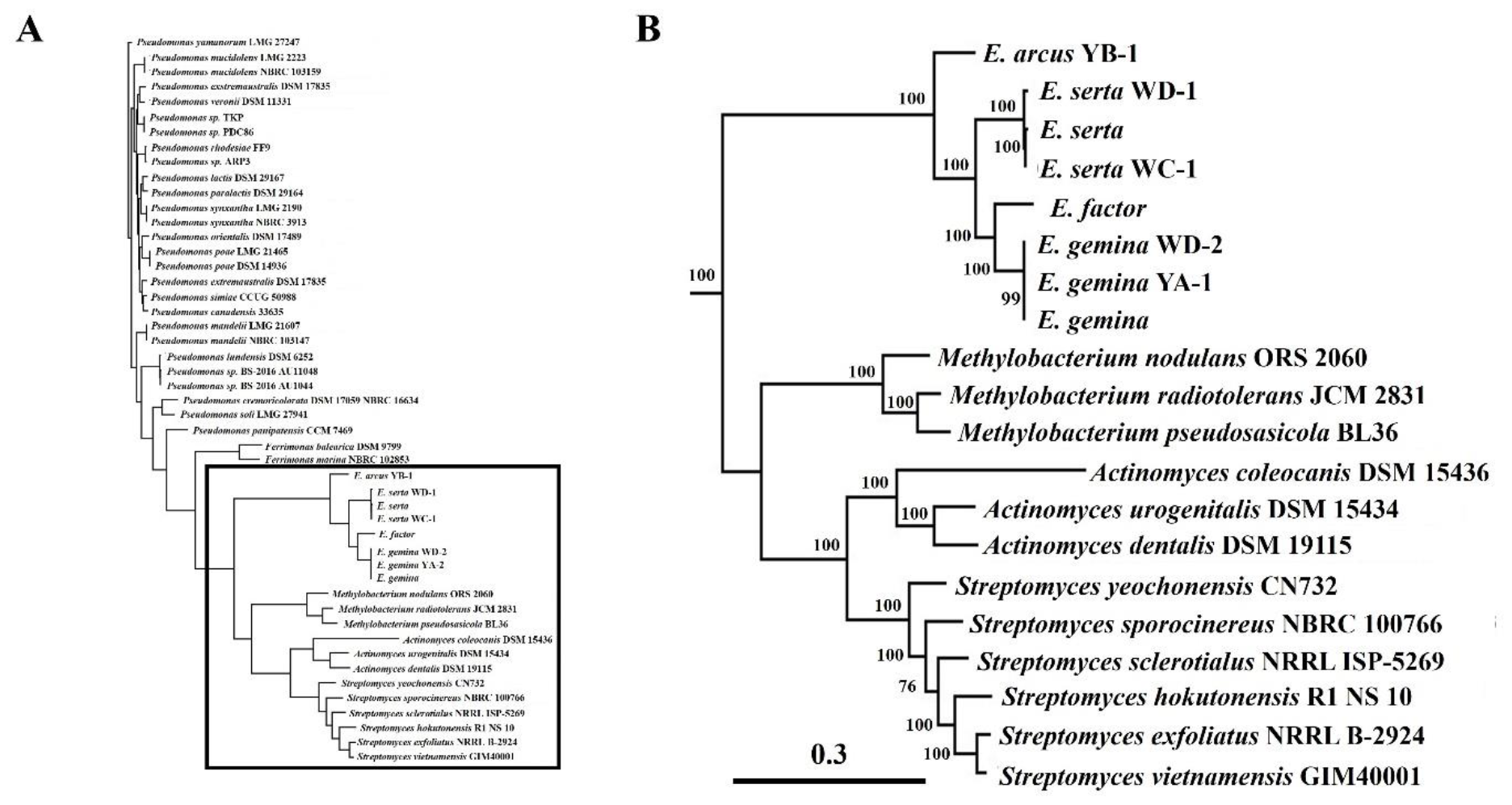
Genome phylogenetic tree of Entotheonella phylotypes. The phylogenetic tree was inferred based on ANI values and common single-copy genes for each genome using the de novo tree mode of autoMLST. Scale bars indicate gene distances. (**A**) The overall phylogenetic tree includes 50 genomes in the query and reference, with reference encompassing *Pseudomonas, Ferrimonas, Bradyrhizobium, Methylobacterium, Actinomyces*, and *Streptomyces* bacteria. (**B**) The enlarged tree including clades of Entotheonella and Actinomycetes are shown.

### Biosynthetic Gene Clusters in Entotheonella Phylotypes

Previously, biosynthetic gene clusters (BGCs) for *Theonella*-derived secondary metabolites have been identified within Entotheonella phylotypes including onnamide (*onn*), keramamide (*ker*), nazumamide *(naz*), cyclotheoneamide (*cth*), konbamide (*kon*), and polytheonamide (*poy*) from *Ca*. E. factor (Wilson et al. 2014); theonellamide (*tna*) (Mori et al. 2018) and misakinolide (*mis*) (Ueoka et al. 2015) from *Ca*. E. serta. To identify putative biosynthetic gene clusters, the genomes were comprehensively analyzed using antiSMASH, an annotation tool for secondary metabolite BGCs (Blin et al. 2023) (Table 2, Table S2). In the genome of YA-1 and YB-1, BGCs related to TSY-derived metabolites, such as *onn, ker, naz, cth, kon*, and *poy*, were identified, whereas BGCs related to TSW-derived metabolites were absent. The genome of WC-1 contained only BGCs for TSW-derived metabolites. In contrast, both WD-1 and WD-2 genomes possess BGCs for TSY-derived (including *ker, naz, cth, kon*, and *poy*) except for *onn*, as well as TSW-derived (*mis* and *tna*) metabolites. The discovery of the *poy* genes in the metagenomes of TSW-derived WD-1 and WD-2 suggests their widespread presence across various Entotheonella variants and sponge hosts, regardless of TSY and TSW origin. This is a noteworthy finding, since the *poy* BGC, initially identified on the plasmid of *Ca*. E. factor, had not been previously reported in TSW-derived samples.

**Table 2.**
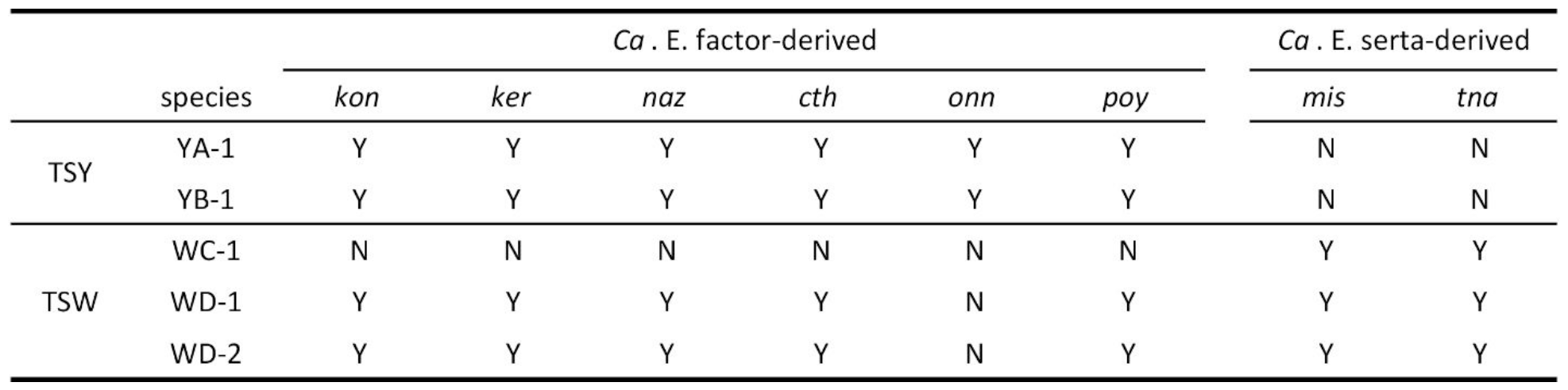
The presence/absence of previously known Entotheonella-derived BGCs in the Entotheonella phylotypes.

Our research also extended beyond those previously known BGCs in *Theonella* phylotypes. In a phylogenetic analysis, YA-1 and WD-2 showed a close relationship with *Ca*. E. gemina. Among the nine BGCs found in the genomic DNA of YA-1, WD-2, and *Ca*. E. gemina, four clusters (BGC-1, 3, 6, and 7) were common to all three, while three others (BGC-2, 4, and 8) were shared between YA-1 and *Ca*. E. gemina (Fig. 4). BGC-4 is the type III PKS cluster responsible for producing the phenolic lipid polyketides. In a previous study, this cluster was shared between *Ca*. E. factor and *Ca*. E. serta (Reiter S et al. 2020). However, we could not detect the BGC in the WD-2 genome. Metabolites derived from the other BGCs remain unknown. These results suggest the presence of certain Entotheonella phylotype is independent of TSY and TSW chemotypes. The genome of YB-1 contains fewer BGCs for secondary metabolites (85 kb) with 65 kb of these BGCs being unique to YB-1. Additionally, an overlap of 20 kb in BGCs is observed in *Ca*. E. factor.

**Fig. 4.**
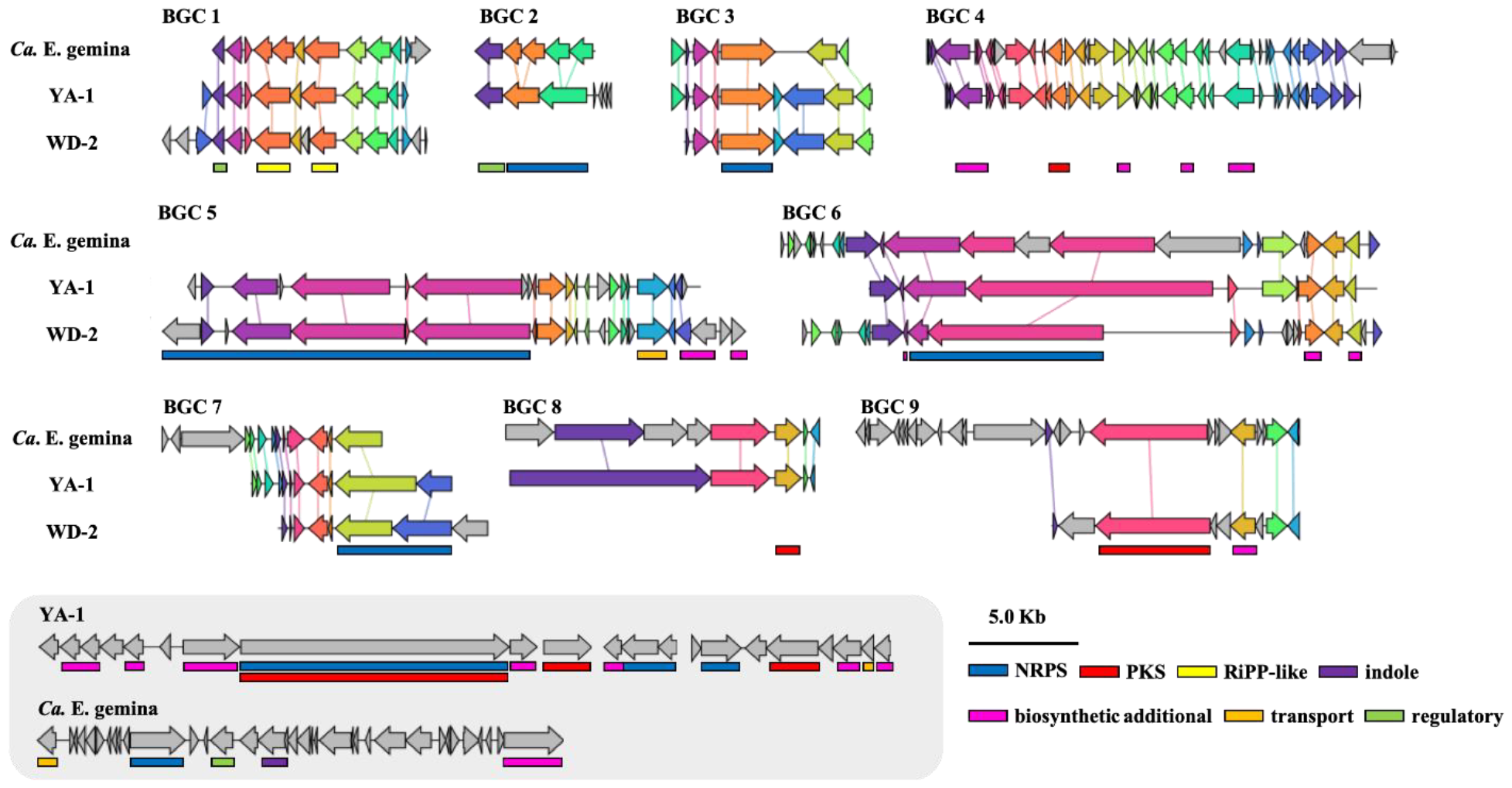
Comparison of BGCs in closely related Entotheonella variants BGC regions in YA-1, WD-2 and *Ca*. E.gemina were aligned and visualized using Clinker. Similar BGCs were grouped and color-coded according to their genes. The core genes of secondary metabolites, as annotated in antiSMASH, are also displayed below the BGCs. The BGCs highlighted in gray indicate non-shared regions among the genomes, suggesting that WD-2 does not possess unique BGCs in comparison to the other two variants, YA-1 and *Ca*. E. gemina.

## Discussion

Numerous secondary metabolites have been discovered in *Theonella* sponges, with the marine sponge *Theonella swinhoei* emerging as a particularly promising source of biologically active substances. Although the biosynthetic pathways and microorganisms responsible for producing many of sponge-derived natural products are not fully understood, the symbionts of *Theonella* sponges are a primary research focus. Previous research identified three Entotheonella phylotypes, *Ca*. E. factor and *Ca*. E. gemina from TSY, and *Ca*. E. serta from TSW, which are known for producing sponge-derived natural products. In our ongoing research, we observed that Entotheonella in both TSY and TSW exhibit several morphologies. Consequently, we became interested in determining the types of BGCs which these Entotheonella possess.

To investigate this, we employed density gradient centrifugation and successfully obtained eight distinct filamentous bacteria-enriched fractions from TSY and TSW. The initial separation using Percoll removed other unicellular bacteria, yielding a single fraction enriched with filamentous bacteria. These bacteria were further separated using an IDX gradient, which enabled the separation of Entotheonella with unique morphologies, as shown in Figures S1 and S2. During density gradient centrifugation with IDX, the addition of HEPES enhanced the separation of Entotheonella.

Subsequent metagenomic and bioinformatic analyses revealed the presence of seven distinct Entotheonella genomes. The Entotheonella genomes we obtained are comparable in quality to those of known Entotheonella, with similar genome size, completeness, and contamination levels (Table 1). The genome size obtained in this study were around 7.0-8.3 Mbp. Although Entotheonella genomes are among the largest known for prokaryotes, an accurate determination of their genome size will require complete genome analyses. Our metagenome analyses led to the discovery of a novel phylotype, *Ca*. E. arcus YB-1, identified through comprehensive ANI and autoMLST analyses (Fig. 2, Fig. 3). Interestingly, despite WC-1, WC-2, and WC-3 being distinctly separated by density gradient centrifugation, our genomic analysis revealed that their genetic profiles are similar to that of *Ca*. E. serta (Fig.2, Fig. S3). YA-1 and WD-2, derived from the different chemotypes TSY and TSW respectively (Fig. 2), exhibit similar morphology, and their genomes are closely related to that of *Ca*. E. gemina. These findings suggest that the diverse morphologies of Entotheonella members do not always directly correlate with phylotypes differentiation. The BGCs in Entotheonella obtained from this study also showed more diversity than expected. WD-1 and WD-2 from TSW possess BGCs for TSY-derived metabolites, such as *kon, ker, naz, cth*, and *poy*, while WC-1 does not harbor them (Table 2). Previous research suggested that *onn* and *poy* genes were detected in the plasmid-derived contigs. We hypothesize that the plasmid was transferred and shared among different Entotheonella via horizontal gene transfer. However, in LC/MS analyses of the fraction Fra7 containing WD-1 and Fra8 containing WD-2, no corresponding metabolites were observed. This could be due to the high cell membrane permeability of polytheonamide leading to their dispersion into the solvent during processing.

This study enhances our understanding of the genomic diversity of Entotheonella, suggesting that *Theonella* sponges host more complex bacterial communities than previously recognized. This complexity may significantly contribute to the chemical defense of these sponges. However, our focus remained primarily on Entotheonella, a dominant group in these sponges. Further comprehensive single-cell genomics studies of Entotheonella in *Theonella* or other lithistid sponges, such as those in the *Discodermia* genus, might reveal even greater diversity of Entotheonella, leading to discovery of untapped bioresources.

## Credit author statement

J.P., H.T., S.M., and K.T. conceived the project and designed the experiments. S.Y., Y.K., A.N., and S.M. performed the experiments. S.Y., S.M., and K.T. wrote the manuscript. All authors reviewed the manuscript and approved the final version to be published.

## Declaration of competing interest

The authors declare no competing financial interest.

## Supporting Information Available

Additional figures and tables are available free of charge via the Internet at ^***^

## Acknowledgements

This work was partially supported by JSPS KAKENHI (grant nos. 17H06403 and 22H02438), the Gordon and Betty Moore Foundation (#9204, https://doi.org/10.37807/GBMF9204), and the Swiss National Science Foundation (SNSF; 205320_185077 and 205320_219638).

## Notes

### Competing Interest Statement

The authors have declared no competing interest.

