## Supplementary file for "Metagenomic Insights Reveal Unrecognized Diversity of Entotheonella in Japanese *Theonella* Sponges"

*Corresponding author

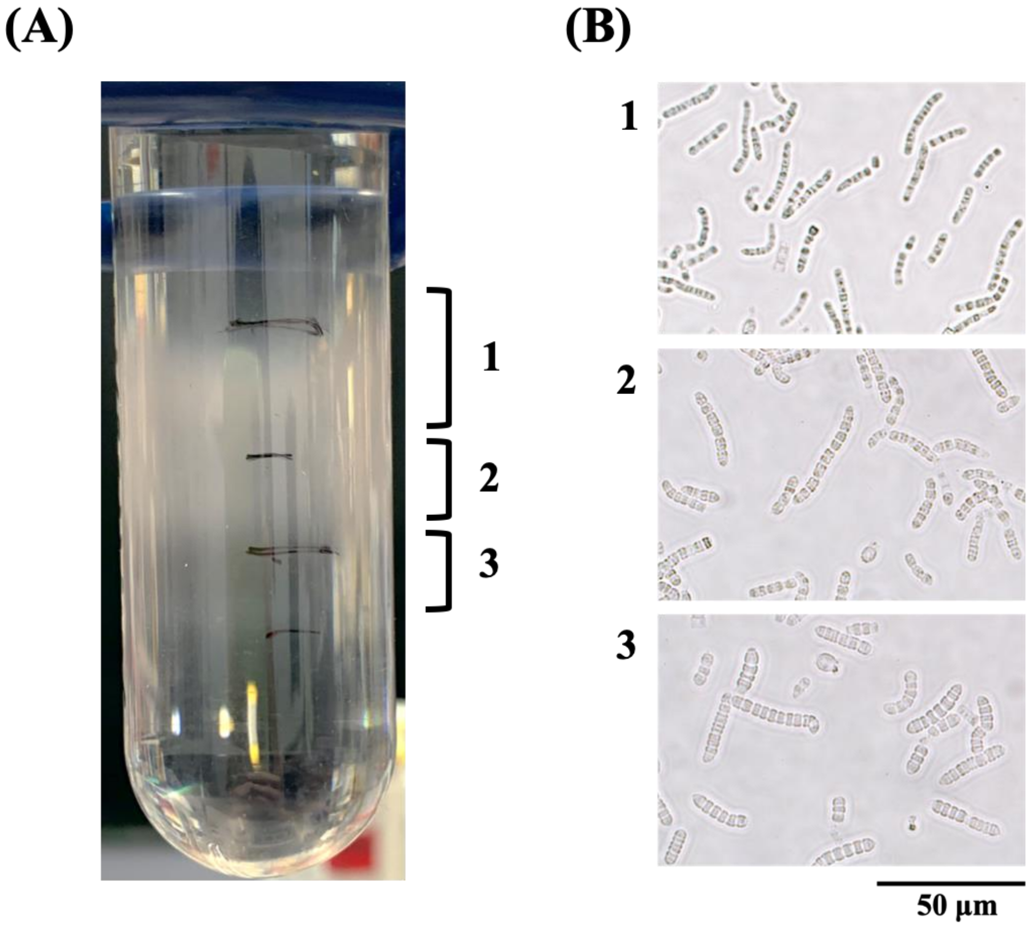


**Figure S1** Results of the separation of filamentous bacteria, enriched through density gradient centrifugation. (A) Tubes showing the fraction after the density gradient centrifugation. (B) Micrographs showing the filamentous bacteria isolated from each fraction.

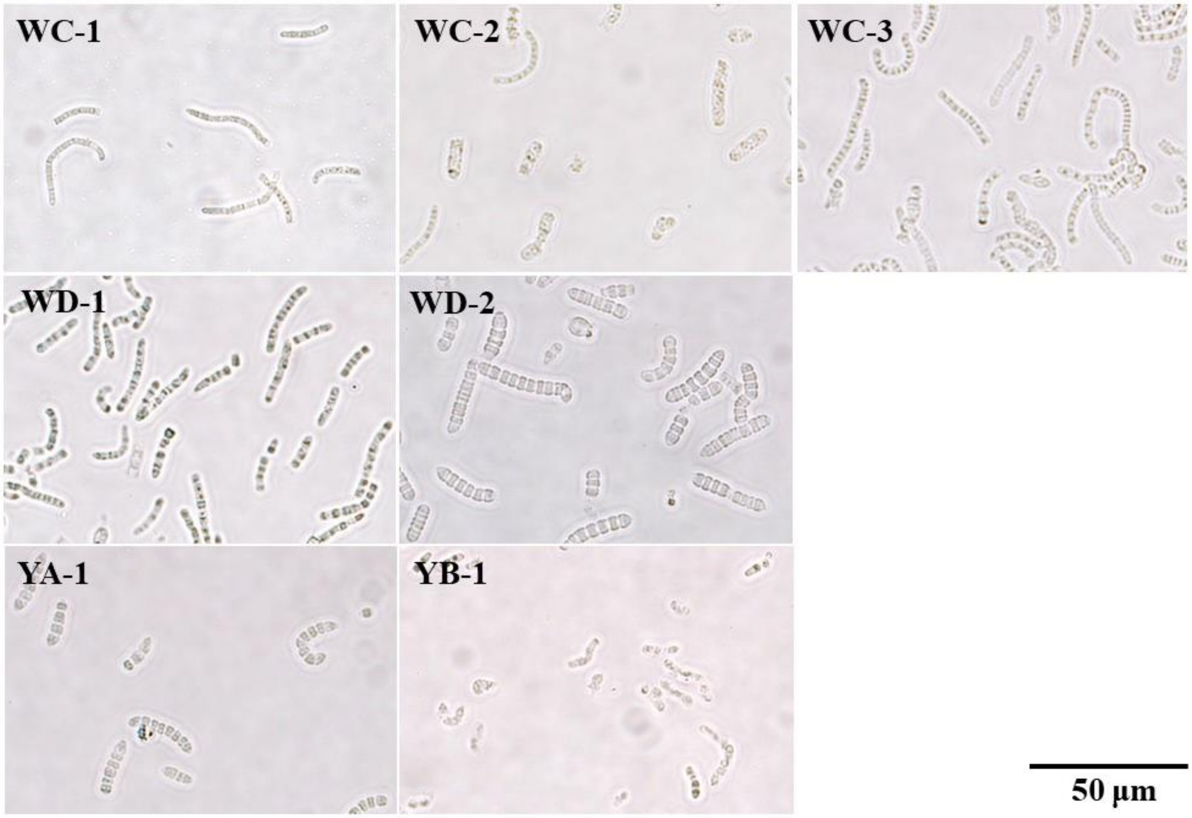


**Figure S2** Micrographs showing Entotheonella-enriched fractions: WC-1, WC-2, and WC-3 from *Theonella swinhoei* (chemotype WC); WD-1 and WD-2 from chemotype WD; YA-1 from chemotype YA; YB-1 from chemotype YB.

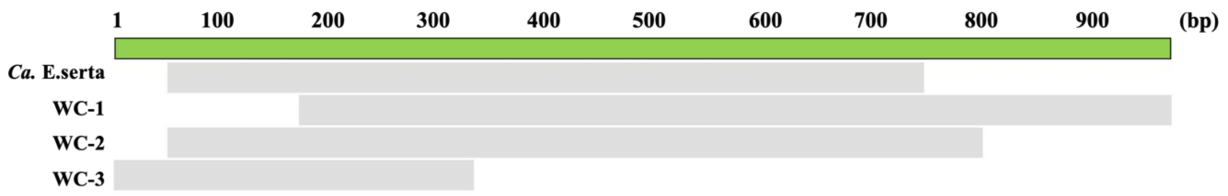


**Figure S3** Alignment results of 16S rRNA gene sequences obtained from *Ca.* E.serta and WC-1, WC-2, and WC-3.

**Table S1.** List of single-copy genes used in the phylogenetic tree by autoMLST.

| NCBI HMM accession | Name | Proposed protein function | Description |
|---|---|---|---|
| TIGR02673 | FtsE | Cellular processes | cell division ATP-binding protein FtsE |
| TIGR01951 | nusB | Transcription | transcription antitermination factor NusB |
| TIGR00033 | aroC | Amino acid biosynthesis | chorismate synthase |
| TIGR00174 | miaA | Protein synthesis | tRNA dimethylallyltransferase |
| TIGR03723 | T6A_TsaD_YgjD | Protein synthesis | tRNA threonylcarbamoyl adenosine modification protein TsaD |
| TIGR01083 | nth | DNA metabolism | endonuclease III |
| TIGR03725 | T6A_YeaZ | Protein synthesis | tRNA threonylcarbamoyl adenosine modification protein YeaZ |
| TIGR00090 | rsfS_iojap_ybeB | Protein synthesis | ribosome silencing factor |
| TIGR01164 | rplP_bact | Protein synthesis | ribosomal protein uL16 |
| TIGR01067 | rplN_bact | Protein synthesis | ribosomal protein uL14 |
| Ribosomal_S8 | PF00410.15 | Unclassified | Ribosomal protein S8 |
| Ribosomal_S9 | PF00380.15 | Unclassified | Ribosomal protein S9/S16 |
| TIGR00061 | L21 | Protein synthesis | ribosomal protein bL21 |
| TIGR00060 | L18_bact | Protein synthesis | ribosomal protein uL18 |
| TIGR00059 | L17 | Protein synthesis | ribosomal protein bL17 |
| TIGR00414 | serS | Protein synthesis | serine--tRNA ligase |
| TIGR00459 | aspS_bact | Protein synthesis | aspartate--tRNA ligase |
| TIGR00755 | ksgA | Protein synthesis | ribosomal RNA small subunit methyltransferase A |
| TIGR00419 | tim | Energy metabolism | triose-phosphate isomerase |
| TIGR00008 | infA | Protein synthesis | translation initiation factor IF-1 |
| TIGR03635 | uS17_bact | Protein synthesis | ribosomal protein uS17 |
| TIGR02273 | 16S_RimM | Transcription | 16S rRNA processing protein RimM |
| TIGR03631 | uS13_bact | Protein synthesis | ribosomal protein uS13 |
| TIGR03632 | uS11_bact | Protein synthesis | ribosomal protein uS11 |
| TIGR00810 | secG | Protein fate | preprotein translocase, SecG subunit |
| TIGR01079 | rplX_bact | Protein synthesis | ribosomal protein uL24 |
| TIGR01011 | rpsB_bact | Protein synthesis | ribosomal protein uS2 |

**Table S2.** Summary of biosynthetic gene cluster for secondary metabolites.

| | ***E.factor*** | ***E.gemina*** | **YA-1** | **YB-1** | ***E.serta*** | **WC-1** | **WD-1** | **WD-3** |
|---|---|---|---|---|---|---|---|---|
| **Total size (Kb)** | 565 | 173 | 182 | 85 | 352 | 381 | 262 | 108 |
| **Number of contigs** | 23 | 10 | 13 | 15 | 28 | 18 | 27 | 8 |
| **GC (%)** | 53.2 | 52.1 | 57.2 | 57 | 54.9 | 54.9 | 56.7 | 56.7 |
| **Number of N bases** | 31,690 | 14,409 | 519 | 411 | 1,287 | 553 | 682 | 0 |

**Table S3.** Biosynthetic domains and enzymes for secondary metabolites in YB-1 compared to previously reported *Entotheonella* variants.

| | ***Ca.* E. factor** | ***Ca.* E. serta** | **YB-1** |
|---|---|---|---|
| **NRPS** | | | |
| Adenylation | 40 | 21 | 15 |
| Condensation | 39 | 30 | 13 |
| Peptidyl-carrier protein | 44 | 27 | 15 |
| Thioesterase | 4 | 4 | 2 |
| Other NRPS domain | 13 | 4 | 4 |
| **PKS** | | | |
| Ketosynthase | 23 | 24 | 1 |
| Acyltransferase | 11 | 8 | 1 |
| Ketoreductase | 12 | 18 | 0 |
| Dehydratase | 8 | 10 | 1 |
| Acyl-carrier protein | 26 | 26 | 0 |
| Thioesterase | 1 | 2 | 0 |
| Other PKS domain | 15 | 17 | 0 |
| *trans*-AT docking | 10 | 16 | 0 |
| Type Ⅲ PKS system | 1 | 3 | 2 |
| **Other second metabolites** | 19 | 18 | 14 |
| **Total** | 266 | 228 | 68 |
